# A Functional Atlas of the Tardigrade Resistome Reveals a Diverse Molecular Toolkit for Extremotolerance

**DOI:** 10.1101/2025.06.10.658754

**Authors:** Takumi Ito, Kohei Ota, Hidekazu Hishinuma, Hideyuki Shimizu

## Abstract

How organisms survive environmental extremes that push life to its physical limits is a fundamental question in biology. Tardigrades are a paradigm for this resilience, entering a state of suspended animation called cryptobiosis to withstand near-complete desiccation and intense radiation. While a few key effectors, including the DNA-shielding protein Dsup and various tardigrade-specific intrinsically disordered proteins (TDPs) have been identified, they constitute only a fraction of a vast, uncharacterized proteome, fundamentally limiting a systems-level understanding of this remarkable biology. Here we overcome this limitation by developing AEGIS, an AI-driven engine that systematically discovers and prioritizes novel ‘guardian’ proteins constituting the tardigrade molecular shield. Applying AEGIS to the tardigrade proteome, we construct the first functional atlas of the tardigrade ‘resistome’, revealing hundreds of novel protein families organized into a diverse and compartmentalized molecular toolkit. This atlas pinpoints new Dsup-like nuclear proteins, suggesting multi-layered genome protection, and a rich cohort of cytoplasmic proteins poised to form protective biological glasses. Our work provides a foundational resource for understanding the evolution of extremotolerance and serves as a blueprint for engineering novel biomolecules with transformative biotechnological potential.

## Introduction

How organisms survive environmental extremes that push life to its physical limits is a fundamental question in biology. Tardigrades, microscopic invertebrates colloquially known as water bears, are renowned for their extraordinary ability to tolerate a range of environmental extremes that are lethal to most other life forms. Through a reversible metabolic suspension known as cryptobiosis—most notably anhydrobiosis (‘life without water’)—many tardigrade species can withstand near-complete desiccation, extreme temperatures, high pressure, vacuum, and intense radiation^1^. This remarkable resilience makes tardigrades a frontier model system for studying the molecular underpinnings of stress tolerance and the fundamental limits of animal life.

The molecular basis for this resilience has long been a subject of intense research. While hypotheses initially centered on common protectants like the disaccharide trehalose^2,3^ and Late Embryogenesis Abundant (LEA) proteins, which are crucial in many anhydrobiotic organisms^4,5^, subsequent studies revealed that many robust tardigrade species accumulate little to no trehalose, suggesting the existence of novel, protein-centric protective mechanisms^6^. The advent of modern genomics has indeed led to the landmark discovery of several families of tardigrade-unique proteins. Prominent among these are the Cytoplasmic Abundant Heat Soluble (CAHS) proteins, which are critical for desiccation tolerance by forming vitrifying biological glasses that restrict molecular motion^6,7^, and the Secreted- and Mitochondrial-(SAHS and MAHS) proteins that protect the extracellular space and mitochondria, respectively^8,9^. Another key discovery is the Damage suppressor (Dsup) protein, a novel DNA-associating protein that confers exceptional radiotolerance^10–12^ by physically shielding from hydroxyl radicals. More recently, other effectors such as the DNA repair-enhancing protein TDR1^13^ and the phase-separating protein TRID1^14^ have further expanded this known toolkit. A unifying feature of these ‘guardian’ proteins is that many are intrinsically disordered proteins (IDPs). Unlike globular proteins, IDPs exist as dynamic conformational ensembles, a structural plasticity that allows them to engage in diverse protective functions^15^. Therefore, these tolerant proteins are called tardigrade disordered proteins / tardigrade-specific intrinsically disordered proteins (TDPs)^16^. The demonstrated ability of these proteins to confer stress tolerance in heterologous systems, including human cells, underscores their immense potential for biomedical and biotechnological applications^17,18^.

Yet, these landmark discoveries likely constitute only a small fraction of a much larger, unexplored molecular toolkit^19^. Genomic and transcriptomic surveys have uncovered a vast number of lineage-specific genes in tardigrades, the majority of which are unannotated open reading frames (ORFs), representing a large, uncharted proteomic landscape^20,21^. Systematically exploring these unknown proteins with traditional experimental or homology-based computational methods presents a significant throughput challenge, especially given the prevalence of rapidly evolving IDPs that defy conventional sequence-based functional inference^22^. While the recent revolution in computational biology, driven by powerful machine learning (ML) techniques and large-scale protein language models (PLMs), offers an unprecedented opportunity to navigate this complexity^23–26^, a dedicated strategy to systematically mine the tardigrade resistome has been lacking.

Here, we overcome this bottleneck by developing AEGIS (AI-driven Engine for Guardian IDPs in Stress-tolerance), a computational framework designed to systematically discover the ‘guardian’ proteins that constitute the molecular shield against extreme stress (Fig. 1). AEGIS first curates a comprehensive tardigrade-specific proteome from public databases through a rigorous filtering pipeline that removes contaminants and conserved orthologs. It then employs an ensemble of fine-tuned protein language models to predict a ‘resistance potential’ score (AEGIS score) for every candidate protein and to map their intrinsically disordered regions (IDRs) at residue-level resolution, enabling a data-driven prioritization of potential novel protective effectors. Applying AEGIS, we have constructed the first extensive functional atlas of the tardigrade resistome, identifying and characterizing hundreds of novel candidate guardian protein families. Our analysis reveals a diverse and compartmentalized molecular toolkit, including new Dsup-like nuclear proteins and cytoplasmic proteins poised to form protective biological glasses. This work provides a foundational resource to accelerate the experimental investigation of tardigrade biology, offering fundamental insights into the diverse molecular strategies of life in extreme environments and opening new avenues for bio-inspired engineering.

**Figure 1.**
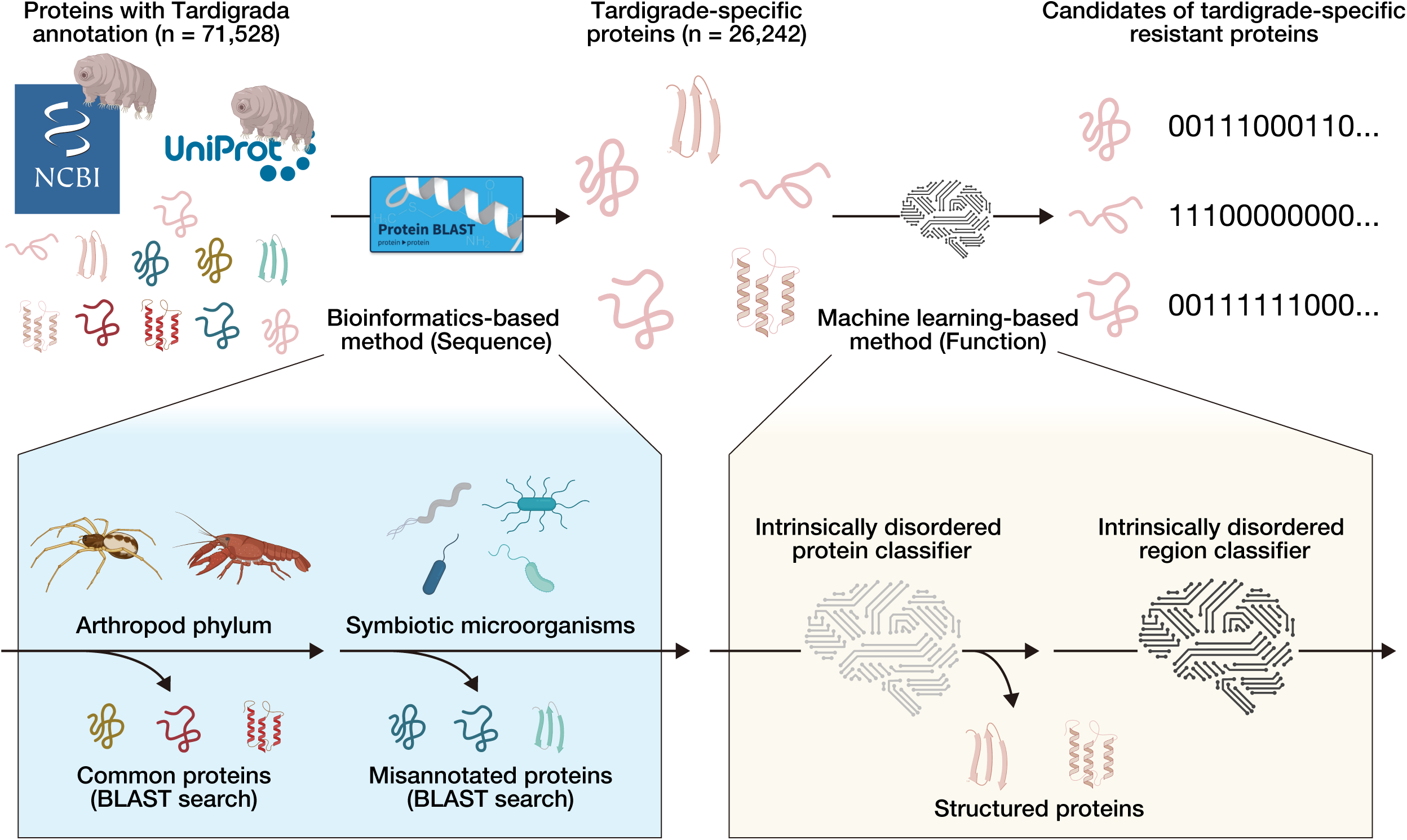
Schematic of the AEGIS Framework for the Systematic Discovery of Tardigrade Guardian Proteins. Schematic representation of the comprehensive computational pipeline, AEGIS (AI-driven Engine for Guardian IDPs in Stress-tolerance), designed for the systematic discovery of novel resistance effectors. The workflow consists of two primary stages. First, in the sequence-based curation stage (left), a raw dataset of all available tardigrade-annotated proteins from public databases (e.g., NCBI, UniProt) is systematically filtered. This step utilizes homology searches (Protein BLAST) to remove conserved proteins shared with other phyla (e.g., Arthropoda) and potential contaminants from symbiotic microorganisms, thereby yielding a high-confidence tardigrade-specific proteome. Second, in the machine learning-based functional prediction stage (right), this curated proteome is processed by a series of classifiers. A protein-level classifier first identifies candidate intrinsically disordered proteins (IDPs), a key feature of many stress-related effectors. A subsequent residue-level classifier then provides fine-grained annotation by mapping the precise locations of intrinsically disordered regions (IDRs). This dual-stage framework enables the high-throughput discovery and prioritization of novel candidate resistance proteins from complex genomic and proteomic data.

## Results

### A Systematic Proteome-wide Survey Reveals a Vast Repertoire of Novel Tardigrade-Specific Protein Families

To systematically identify proteins unique to the phylum Tardigrada, we first sought to establish a comprehensive and high-confidence dataset of their protein repertoire. While landmark studies have identified key stress-resistance proteins such as TDPs^6,8^, these were often discovered through targeted experimental approaches, leaving the vast majority of the tardigrade proteome uncharacterized. We hypothesized that a systematic, proteome-wide survey would uncover a large, previously hidden collection of tardigrade-specific protein families. To test this hypothesis and to overcome the significant challenge of microbial contamination in sequencing microscopic animals^27,28^, we implemented a rigorous bioinformatic pipeline (Fig. 1 left). We began by compiling an initial dataset of 71,528 protein sequences with ‘Tardigrada’ annotation from public databases such as NCBI and UniProt, covering key species including the strong anhydrobiote *Ramazzottius varieornatus* and the model species *Hypsibius exemplaris* (Supplementary Table S1). We then performed exhaustive homology searches using BLASTp against the NCBI non-redundant (nr) database to systematically remove proteins with significant homology to orthologs in related phyla, such as Arthropoda, as well as to screen out sequences likely originating from common symbiotic or environmental microorganisms. To further organize this curated dataset and identify putative protein families, we applied the sequence clustering tool CD-HIT^29^ to enable an unbiased exploration of the tardigrade proteome without relying on existing annotations.

Our bioinformatics pipeline successfully defined a high-confidence tardigrade-specific proteome and revealed its family organization. The multi-stage filtering process reduced the initial set of 71,528 proteins to a final, high-confidence dataset of 26,242 tardigrade-specific protein sequences (Fig. 2a). Analysis of this curated proteome revealed that it was predominantly composed of sequences from *Ramazzottius varieornatus* (7,565 sequences), *Hypsibius exemplaris* (5,373 sequences), and *Paramacrobiotus metropolitanus* (4,201 sequences) (Fig. 2b).

**Figure 2.**
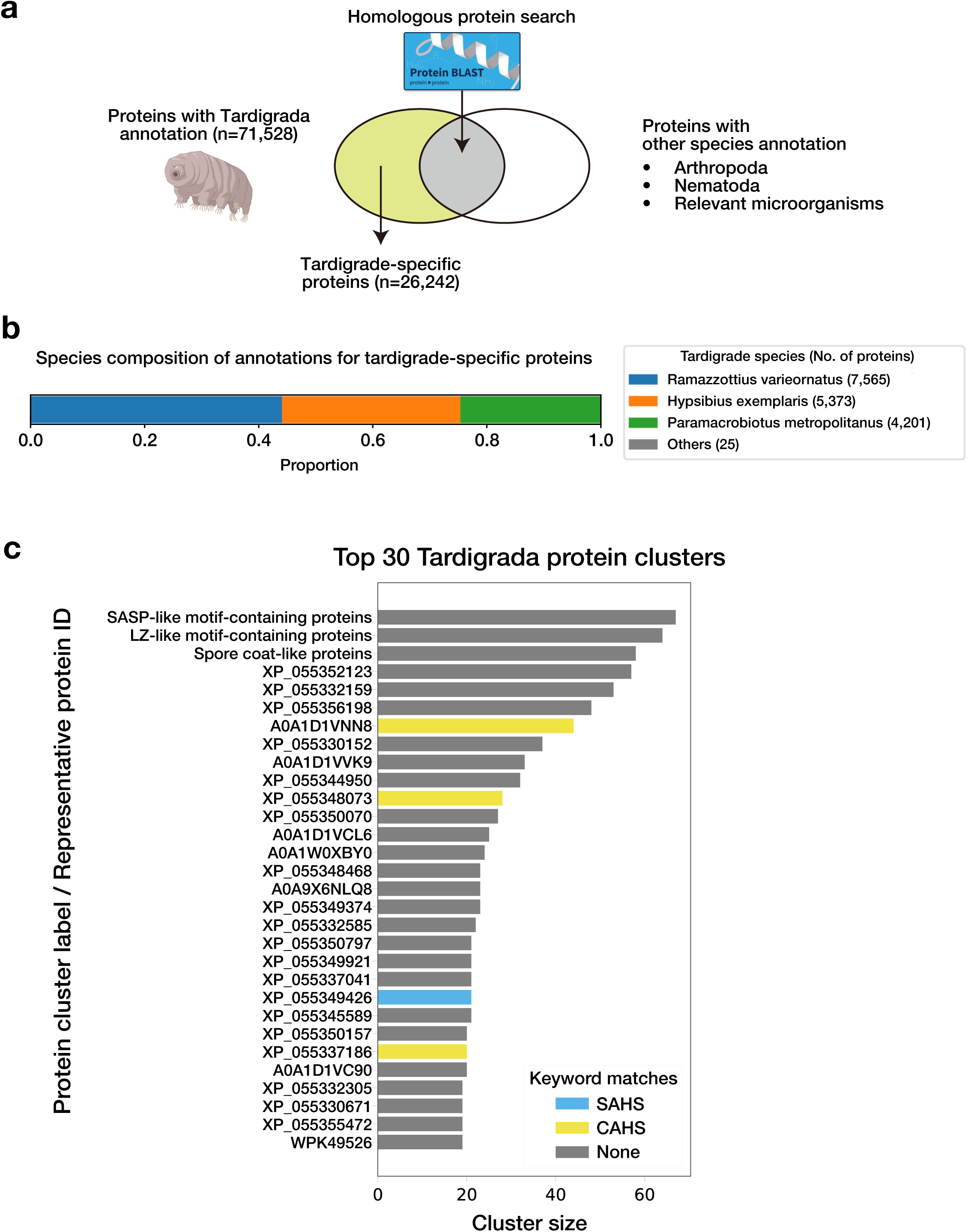
Systematic Curation of the Tardigrade-Specific Proteome Reveals Numerous Novel Protein Families. **(a)** Venn diagram illustrating the outcome of the homology-based filtering pipeline. An initial set of 71,528 annotated tardigrade proteins was curated to a high-confidence dataset of 26,242 tardigrade-specific proteins after removing sequences with significant homology to non-tardigrade organisms. **(b)** Species composition of the curated tardigrade-specific proteome. The dataset is primarily composed of proteins from the strong anhydrobiote *Ramazzottius varieornatus* (7,565)*, Paramacrobiotus metropolitanus* (5,373), and the established model species *Hypsibius exemplaris* (4,201), three of the first tardigrades to have their genomes sequenced. **(c)** The 30 largest protein clusters from the tardigrade-specific dataset, ranked by size. The successful grouping of known CAHS (yellow) and SAHS (blue) proteins into discrete clusters validates this clustering approach. The majority of the largest clusters are composed of unannotated proteins (gray). Motif analysis of the top three unannotated families suggests putative functions: the largest contains a motif similar to that of DNA-protecting Small, Acid-Soluble Spore Proteins (SASPs); the second contains a Leucine Zipper-like (LZ-like) motif; and the third contains spore coat-like proteins.

Next, to organize this curated proteome into putative protein families and to enable the discovery of novel ones, we performed sequence-based clustering. We hypothesized that an unbiased clustering approach would not only recapitulate the organization of known tardigrade protein families but also reveal novel, large families that have been overlooked by targeted biochemical methods. For this purpose, we applied the sequence clustering tool CD-HIT to the 26,242 tardigrade-specific proteins, grouping them into putative families based on sequence similarity. We identified 17,164 putative protein families of varying sizes (Supplementary Table S2). Notably, when we examined the 30 largest clusters, we found that our pipeline successfully and correctly grouped known members of the Cytoplasmic Abundant Heat Soluble (CAHS) and Secretory Abundant Heat Soluble (SAHS) protein families into their own distinct clusters (Fig. 2c). For instance, the cluster represented by the sequence XP_055349426 was identified as a major SAHS family group, while the cluster represented by A0A1D1VNN8 corresponded to a CAHS family.

While this validated our approach, it also revealed that the largest clusters, representing a significant portion of the tardigrade-specific proteome, remained entirely unannotated (Fig. 2c, gray bars). These families lacked any known domains or homology to proteins outside the Tardigrada phylum. To gain the first functional insights into these prominent ‘dark matter’ protein families, we next sought to identify conserved sequence motifs within each cluster and infer their potential functions based on similarity to known motifs.

We applied the MEME suite^30^ to the three largest novel protein clusters, which revealed a unique and statistically significant conserved motif for each family. To infer their functions, these novel motifs were then compared against the PROSITE database of known motifs using TOMTOM^31^. This analysis yielded functional hypotheses (Fig. 2c, top labels). The motif from the largest cluster showed significant similarity to the DNA-binding motif of Small, Acid-Soluble Spore Proteins (SASPs), which are known to protect the DNA in bacterial spores from UV damage and other stresses^32^. The motif from the second-largest cluster resembled a leucine zipper-like (LZ-like) motif, suggesting a potential role in dimerization or protein-protein interactions^33^. The third-largest cluster contained proteins that were independently annotated by UniProt as ‘Spore coat-like proteins’, further implicating this family in forming protective structures to safeguard the genome.

These motif-based functional predictions, all pointing towards DNA protection, oligomerization, and structural roles reminiscent of highly resistant bacterial spores, provide the first testable hypotheses for these previously unknown protein families. This suggests that tardigrades may have independently evolved or co-opted a ‘spore-like’ molecular strategy to protect their genome during cryptobiosis. Together, these results establish a comprehensive map of the tardigrade-specific proteome, and the identification of these numerous, large, and functionally intriguing novel protein families offers a rich discovery platform for identifying novel functional effectors through machine learning.

### AEGIS, an AI-driven Framework, Accurately Predicts Stress-Resistance Potential

To functionally interrogate the 26,242 tardigrade-specific proteins and prioritize those likely involved in extremotolerance, we next sought to develop a machine learning framework capable of predicting a protein’s ‘stress-resistance potential’ directly from its primary sequence. Given that many known tardigrade effectors are intrinsically disordered proteins (IDPs)^16^, which are challenging to characterize by traditional homology-based methods^22^, we hypothesized that a specialized deep learning model could capture the subtle sequence features indicative of their function.

We therefore engineered AEGIS (AI-driven Engine for Guardian IDPs in Stress-tolerance), a two-tiered classification framework. First, a protein-level classifier was designed to differentiate between experimentally-validated, disordered proteins like CAHS proteins (positive set) and proteins confirmed to be structured throughout their length (negative set) (Fig. 3a). This classifier was then used to score the entire tardigrade-specific proteome, generating a ranked list of proteins based on their likelihood of being globally disordered. Second, for the top-scoring candidates from this initial screen, we applied a fine-grained, residue-level classifier to precisely map their IDRs. Both classifiers were powered by embeddings from ESM-C, a state-of-the-art PLM, enabling us to capture the nuanced sequence patterns of disorder without reliance on homology^34^. To enhance predictive performance for this unique biological problem, we employed Low-Rank Adaptation (LoRA) to fine-tune the entire embedding framework^35^.

**Figure 3.**
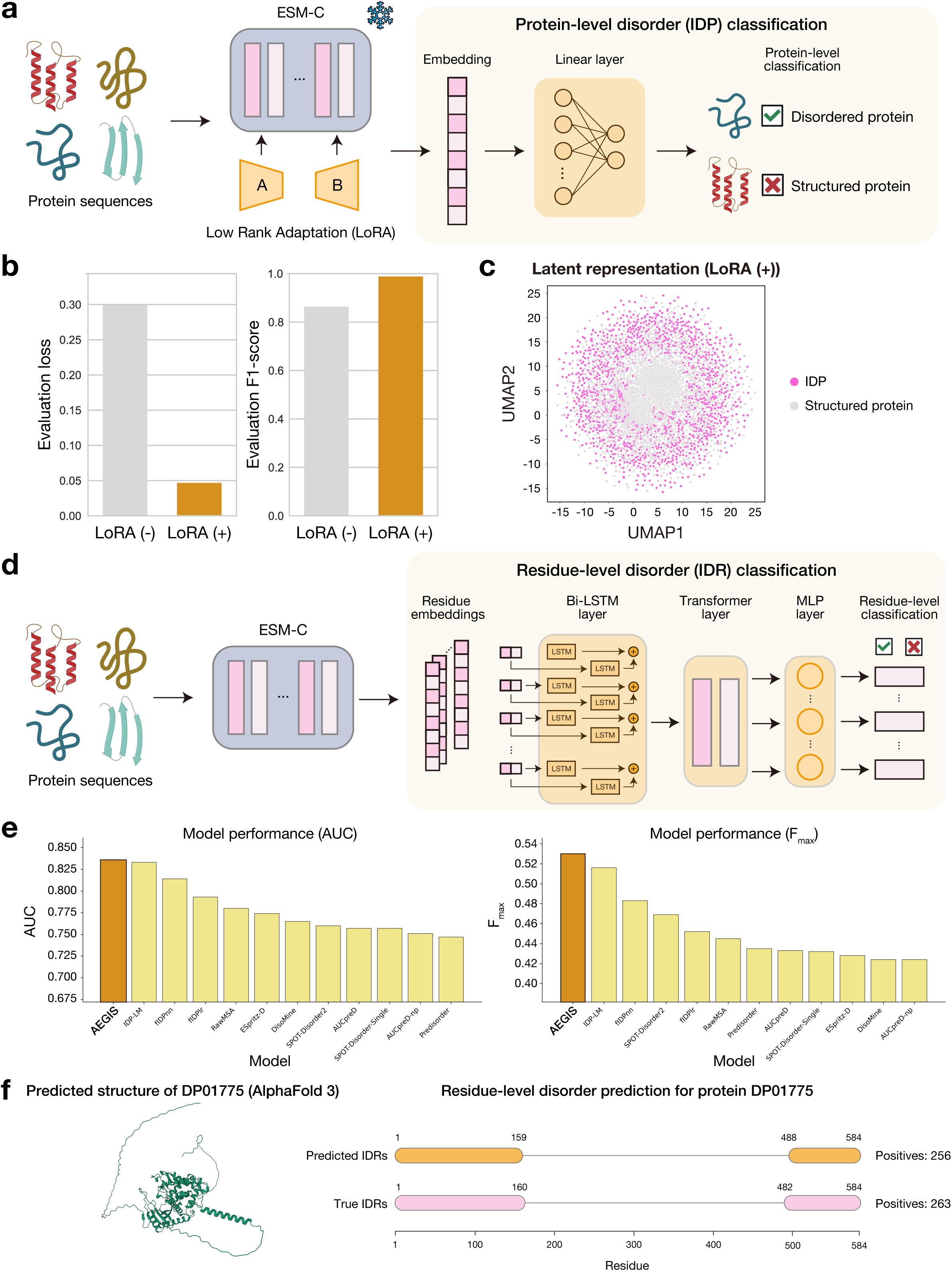
Development and Validation of AEGIS, a State-of-the-Art Protein Disorder Predictor. **(a)** Schematic of the protein-level IDP classifier within the AEGIS framework. This model evaluates the holistic properties of a protein to classify it as either globally disordered or structured. **(b)** Performance of the protein-level classifier with and without Low-Rank Adaptation (LoRA) fine-tuning. The LoRA-tuned model (LoRA (+)) achieves significantly lower evaluation loss and a higher F1-score compared to the baseline model (LoRA (-)), confirming the benefit of the adaptation. **(c)** UMAP visualization of the latent protein embedding space (960 dimensional embedding of CLS token) after LoRA fine-tuning, demonstrating a clear and distinct separation between the intrinsically disordered protein (IDP) and structured protein classes from the training set. **(d)** Schematic of the residue-level IDR prediction workflow. AEGIS processes per-residue embeddings through a Bi-directional Long Short-Term Memory (Bi-LSTM) layer to capture local sequential context, followed by a Transformer encoder layer to model long-range dependencies. The final output is passed through a multilayer perceptron (MLP) to predict disorder for each residue. See Figure S1 and Methods for additional information. **(e)** AEGIS outperforms other state-of-the-art predictors on the independent CAID benchmark dataset. AEGIS ranks first among 12 other methods in both Area Under the Curve (AUC) and maximum F1-score (Fmax). See Table 1 for additional metrics. **(f)** Example residue-level IDR predictions made by AEGIS for protein DP01775 from the held-out CAID test dataset, showing high concordance between predicted and true IDRs. The values on the right of the plot indicate the number of IDR-positive residues.

**Table 1.** Performance comparison of IDR prediction methods.

| Model | AUC | Fmax | MCC | BACC |
| --- | --- | --- | --- | --- |
| **AEGIS (ours)** | **0.836** | **0.530** | **0.427** | **0.763** |
| IDP-LM<sup>36, *</sup> | 0.833 | 0.516 | 0.415 | 0.762 |
| fIDpnn<sup>38, **</sup> | 0.814 | 0.483 | 0.370 | 0.720 |
| fIDPIr<sup>38, **</sup> | 0.793 | 0.452 | 0.330 | 0.693 |
| RawMSA<sup>40, **</sup> | 0.780 | 0.445 | 0.328 | 0.714 |
| ESpritz-D<sup>42, **</sup> | 0.774 | 0.428 | 0.307 | 0.703 |
| DisoMine<sup>37, **</sup> | 0.765 | 0.424 | 0.299 | 0.698 |
| SPOT-Disorder2<sup>39, **</sup> | 0.760 | 0.469 | 0.349 | 0.725 |
| AUCpreD<sup>41, **</sup> | 0.757 | 0.433 | 0.318 | 0.712 |
| SPOT-Disorder-Single<sup>39, **</sup> | 0.757 | 0.432 | 0.315 | 0.710 |
| AUCpreD-np<sup>41, **</sup> | 0.751 | 0.424 | 0.301 | 0.699 |
| Predisorder<sup>43, **</sup> | 0.747 | 0.435 | 0.301 | 0.691 |

\* Values for IDP-LM were obtained from the original IDP-LM paper, reporting results of evaluation on the same CAID dataset.
\*\* Values for other models were obtained from the CAID report.

This fine-tuning process resulted in a marked improvement in model performance. Quantitatively, the fine-tuned model (LoRA (+)) achieved a significantly lower evaluation loss and a higher F1 score compared to the model without fine-tuning (LoRA (-)) (Fig. 3b). The latent representation of the training data from the fine-tuned model showed a clear and distinct separation between the IDP and structured protein classes (Fig. 3c). This demonstrates that our fine-tuning strategy generates a more powerful and context-aware feature representation, enhancing the model’s ability to distinguish between globally disordered and structured proteins accurately. This initial screening provided a high-confidence, prioritized list of putative IDP candidates for further analysis.

To enable the precise mapping of IDRs, we trained a second classifier model that predicts disorder at the residue level (Fig. 3d, Supplementary Figure S1). The classifier was trained using residue level ESM-C embeddings of IDPs recorded in MobiDB. The classifier utilizes both the LSTM and Transformer architecture to capture the intricate local and long-range sequence dependencies which contribute to disorder.

To rigorously validate the overall performance of our final AEGIS framework, we then benchmarked it against a suite of twelve widely used, state-of-the-art IDR predictors^36–43^. This was performed using the standardized Critical Assessment of Intrinsic Disorder (CAID) dataset^44^ to ensure an unbiased comparison. On this independent benchmark, AEGIS demonstrated superior performance, ranking first among all tested methods. Specifically, AEGIS achieved the highest Area Under the Curve (AUC) score of 0.836 and the highest maximum F1-score (Fmax) of 0.530 (Fig. 3e). It also outperformed all other predictors on the Matthews Correlation Coefficient (MCC) and Balanced Accuracy (BACC) metrics (Table 1). Examining the prediction results of a specific protein showed that AEGIS was successful in predicting the IDR of an unseen protein sequence, spanning over 250 residues at the N-terminus and C-terminus (Fig. 3f).

These results firmly establish AEGIS as a new state-of-the-art method for predicting protein disorder. While fine-tuning PLMs using parameter efficient methods such as LoRA is a well-established practice, our work demonstrates its application to the specific challenge of identifying and annotating IDPs. The superior performance of AEGIS, benchmarked against the field’s leading tools, validates our two-tiered framework and its ability to bridge the sequence-function gap for IDPs. This gives us high confidence in its application to the novel tardigrade proteome for discovering new stress-response protein candidates.

### AEGIS-driven Analysis Reveals a Diverse Toolkit of Novel Guardian Proteins

With the validated AEGIS model in hand, we next applied it to the entire 26,242-protein tardigrade-specific proteome to create a systems-level functional map and prioritize novel candidates for in-depth characterization. We used AEGIS to assign a score, representing the predicted stress-resistance potential, to each of the 26,242 tardigrade-specific proteins (Fig. 4a). A UMAP visualization of the protein embedding space, colored by the allocated cluster and the AEGIS score, revealed distinct ‘hot clusters’ of high-scoring proteins across the proteomic landscape (Fig. 4b, left and middle). To validate the biological relevance of this score, we mapped the positions of known TDPs onto this landscape. This analysis revealed a strong correlation between the AEGIS score and the structural nature of these known effectors (Fig. 4b, right). Notably, proteins known to be highly disordered, such as members of the CAHS and Dsup families, were consistently located within the highest-scoring regions (bright yellow). In contrast, TDPs known to adopt more structured globular folds, such as members of the SAHS family^6^, received comparatively lower scores (darker colors). Crucially, we observed several prominent hotspots populated by uncharacterized proteins that received even higher AEGIS scores than the known CAHS and Dsup proteins. This result indicates that the AEGIS score serves as a reliable proxy for identifying highly disordered proteins for tardigrade, strongly suggesting that these particularly high-scoring regions harbor novel, potent TDPs. This provides a powerful, data-driven method to prioritize the most promising candidates for subsequent characterization.

**Figure 4.**
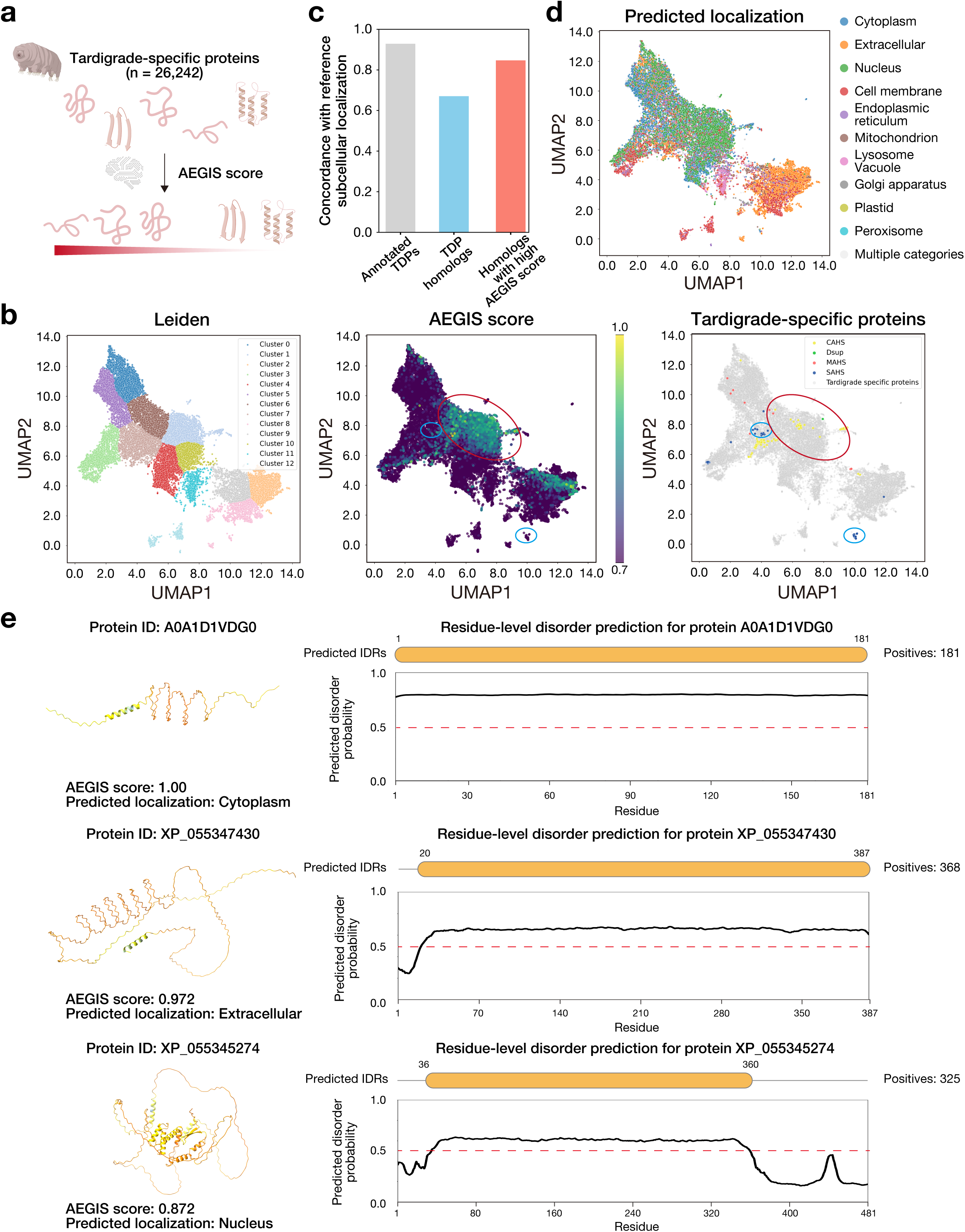
AEGIS-driven Analysis Reveals a Diverse and Compartmentalized Toolkit of Novel Guardian Proteins. **(a)** Schematic showing that the tardigrade-specific proteins filtered in Fig. 2 were scored with AEGIS. **(b)** UMAP visualization of the 26,242 tardigrade-specific proteins, colored by Leiden cluster (left) and AEGIS score (middle). High scores (bright yellow) indicate a high predicted potential for stress-resistance function. Notably, highly disordered effectors like CAHS and Dsup map to the highest-scoring regions (red circle, right), while more structured proteins like SAHS (blue circles) receive comparatively lower scores, validating that the AEGIS score is a proxy for functional disorder. **(c)** AEGIS score enriches for functionally relevant TDP homologs. The bar plot demonstrates the utility of the AEGIS score by comparing the accuracy of the subcellular localization predictor across three protein sets. The left bar (gray) shows that annotated TDPs (e.g., annotated CAHS and SAHS proteins) have high concordance with their expected localizations, validating the predictor’s applicability to tardigrade proteins. The middle bar (blue) reveals that prediction accuracy drops significantly for a broad set of sequence homologs, suggesting contamination of proteins not related to resistance and that sequence similarity alone is insufficient to identify functionally equivalent proteins among rapidly evolving IDPs. The right bar (red) shows that filtering this homolog set for a high AEGIS score restores prediction concordance to the annotated level. **(d)** UMAP visualization of the 26,242 tardigrade-specific proteins, colored by the predicted subcellular localization. Proteins predicted to localize in multiple compartments are categorized into ‘Multiple categories’. **(e)** Representative examples of novel ‘guardian protein’ candidates discovered and prioritized by AEGIS. For each candidate, the predicted 3D structure (AlphaFold 3), AEGIS score, and the residue-level disorder probability track from AEGIS are shown. The values on the right of the plot indicate the number of IDR-positive residues. From top to bottom: A0A1D1VDG0, a top-scoring (1.000) cytoplasmic protein that is almost entirely disordered, reminiscent of the CAHS family; XP_055347430, a high-scoring (0.972) extracellular protein that is largely disordered; and XP_055345274, a nuclear protein (0.872) with a unique, more structured fold, representing a potential new class of DNA-associated effector.

To translate these high-scoring predictions into concrete biological hypotheses, we then performed a multi-faceted *in silico* characterization of the top-ranked candidates, integrating predictions of subcellular localization and 3D structure. The subcellular localization prediction model^45^ made accurate predictions for known TDPs, demonstrating the validity of its usage for tardigrade proteins (Fig. 4c). We demonstrated that the concordance rate decreased when including homologs of annotated TDPs, suggesting contamination of proteins not related to resistance with traditional sequence-based approach. When filtering those homologs with AEGIS score, we observed the improvement in the concordance, indicating the importance of combining our methodology.

The model predicted high-scoring candidates were mainly enriched in the cytoplasm and nucleus, suggesting a functionally compartmentalized defense system (Fig. 4d). To illustrate the power of this integrative approach, we highlight three novel, high-scoring candidates from our atlas (Fig. 4e). Of these, two were selected directly from the top-ranked AEGIS predictions, while the third was drawn from one of the principal sequence-similarity clusters on the basis of its exceptionally high AEGIS score. The first, A0A1D1VDG0 (AEGIS score: 1.00, rank = 1), is a fully disordered protein predicted to be cytoplasmic, strongly reminiscent of the CAHS family of vitrifying agents. The second, XP_055347430 (AEGIS score: 0.972, rank = 3), is a protein predicted to be secreted, with a few structured residues. The third candidate, XP_055345274 (AEGIS score: 0.872) belongs to the fifth-largest cluster of tardigrade-specific proteins (Fig. 2c), which is composed of high-scoring members among which XP_055345274 ranks highest. The sequence harbors a motif similar to SASPs, which protect DNA from UV-induced damage, suggesting that XP_055345274 may define a novel class of DNA-binding effectors. This hypothesis is further bolstered by subcellular localization predictions placing the protein in the nucleus. This integrative analysis, combining a robust functional score with structural and localization data, allowed us to construct a rich, hypothesis-generating catalog of novel tardigrade effector proteins with putative roles spanning DNA protection, cytoplasmic stabilization, and extracellular functions.

These results, taken together, provide strong *in silico* evidence that our ML-prioritized list contains a diverse array of bona fide stress-resistance candidates. This integrative analysis has allowed us to construct a rich, hypothesis-generating catalog of novel tardigrade effector proteins whose varied structural properties and predicted localizations suggest a complex, compartmentalized cellular defense system. This work significantly expands the known molecular repertoire for anhydrobiosis and provides a wealth of high-priority targets for future experimental validation.

## Discussion

In this study, we developed and applied the comprehensive computational framework named AEGIS (AI-driven Engine for Guardian IDPs in Stress-tolerance). By deploying this AI-driven discovery engine, we systematically explored the uncharacterized tardigrade proteome and discovered hundreds of novel ‘guardian Protein’ candidates predicted to possess universal properties of known stress-resistant proteins. Our subsequent structural and functional analyses of these candidates have unveiled a surprisingly diverse and compartmentalized molecular toolkit that tardigrades utilize to survive extreme environments. The significance of this work is threefold. First, it provides the first functional atlas to capture the tardigrade stress-response machinery at a systems level, moving beyond the ‘tip-of-the-iceberg’ view provided by studies of individual protein families. Second, our machine learning approach, which transcends the limitations of homology, presents a new discovery paradigm for functional genomics in non-model organisms that possess many unique genes. Finally, the novel molecular cohort identified here represents a valuable genetic resource for future biotechnology and medical applications, such as the stabilization of biologics and radioprotection.

Our work marks a paradigm shift away from the targeted, often serendipitous discovery of individual protein families towards a systematic, proteome-wide exploration of tardigrade biology. While foundational research in the field have successfully identified key protein families like CAHS, SAHS, and Dsup through labor-intensive experimental approaches such as heat-solubility assays or chromatin fractionation, these methods are inherently limited in scope and throughput. Recent genomic analyses have revealed that the tardigrade genome contains thousands of unannotated, lineage-specific open reading frames, representing a large uncharted territory of tardigrade biology. Our systematic, proteome-wide filtering and clustering approach directly addresses this challenge, moving beyond a candidate-gene focus to a truly exploratory discovery model. Crucially, the biological relevance of our approach was validated by the finding that our unbiased, sequence-driven clustering correctly grouped known CAHS and SAHS proteins into their respective families (Fig. 2c). This provides high confidence that the numerous large, unannotated clusters we identified represent bona fide novel protein families with potential roles in tardigrade-specific biology, thereby providing the first foundational atlas of this unique proteome.

A central technological contribution of this work is the development and validation of AEGIS, a highly accurate predictive engine that deciphers the sequence-to-function grammar of stress-resistance proteins. Predicting protein function *in silico* is a central challenge in computational biology, a challenge particularly acute for the tardigrade proteome, which is rich in IDPs that defy classical homology-based inference due to their rapid sequence evolution. We therefore engineered AEGIS to build upon the transformative power of PLMs, demonstrating that an ensemble of models fine-tuned on domain-specific data yields superior predictive performance (Fig. 3). The high accuracy of AEGIS establishes a robust, data-driven framework to navigate the vast functional landscape of the uncharacterized tardigrade proteome, enabling the efficient prioritization of high-value targets where experimental validation is resource-intensive.

A key advance of our integrative approach is the ability to generate specific, systems-level hypotheses about the organization of the tardigrade defense network. Unlike transcriptomic analyses, which are limited to specific conditions and species, or homology-based searches, which can only identify analogs of known proteins, our strategy has the power to uncover entirely novel classes of effectors. For instance, we identified numerous highly disordered cytoplasmic proteins that are prime candidates for forming protective vitrified glasses or undergoing liquid-liquid phase separation (LLPS) to arrest molecular motion during desiccation, as hypothesized in CAHS proteins. Furthermore, the discovery of novel nuclear-localized, high-scoring families distinct from Dsup suggests that tardigrades employ a multi-layered strategy for genome protection. Collectively, these findings paint a picture of tardigrade anhydrobiosis not as a single mechanism, but as a complex and integrated system of specific defenses tailored to different types of stress and cellular compartments.

This study has several limitations. First, our findings are entirely *in silico*. Although the predictions made by AEGIS are supported by rigorous computational validation, the putative functions of the identified candidate proteins must ultimately be confirmed through experimental approaches. Second, the comprehensiveness of our study is necessarily dependent on the protein sequences currently available in public databases. Consequently, any undiscovered gene products not yet identified or deposited are outside the scope of this analysis. However, our framework is designed to be scalable and can be readily applied as more sequence data become available. Finally, our analysis primarily focuses on primary protein sequences and static structures. It does not account for dynamic regulatory layers such as post-translational modifications or the differential gene expression patterns known to be critical for the anhydrobiotic response. A more complete and dynamic understanding will require the future integration of multi-omics data.

In conclusion, by developing and applying the AI-driven discovery engine AEGIS, this study systematically identifies hundreds of novel ‘guardian protein’ candidates lurking in the largely uncharted tardigrade proteome, elevating our understanding of the molecular basis for their extraordinary resilience to a new, systems-level horizon. The novel molecular cohort identified here represents a valuable genetic resource for realizing next-generation biotechnologies, such as cold-chain-free preservation of vaccines and radioprotection.

## Materials and Methods

### Data Acquisition and Curation of the Tardigrade-Specific Proteome

To construct a comprehensive protein dataset for the phylum Tardigrada, we systematically retrieved all available protein sequences from the NCBI (National Center for Biotechnology Information) GenBank and the UniProt Knowledgebase (UniProtKB) as of June 2025. The initial search query targeted all entries under the NCBI taxonomy ID for Tardigrada (taxid: 42241), yielding a raw dataset of 71,528 protein sequences. This dataset was primarily composed of proteins from the extensively sequenced species *Ramazzottius varieornatus*, *Paramacrobiotus metropolitanus*, and *Hypsibius exemplaris.* These species represent a spectrum of anhydrobiotic capabilities, with *R. varieornatus* and *P. metropolitanus* considered strong anhydrobiotes and *H. exemplaris* a comparatively weak one. This composition provides a valuable comparative framework for investigating the molecular basis of extremotolerance. Prior to filtering, initial quality control was performed, which involved the removal of redundant identical sequences and protein fragments shorter than 50 amino acids to ensure the integrity of the downstream analyses.

To generate a high-confidence set of proteins unique to tardigrades, a rigorous multi-stage filtering pipeline was designed and implemented (Fig. 2a). This pipeline was designed to achieve two key objectives: first, to enrich for proteins uniquely associated with cryptobiosis in tardigrades, and second, to address the significant challenge of microbial and cross-phyla contamination inherent to the sequencing of microscopic animals. To remove orthologs that are broadly conserved with closely related metazoan phyla, we first performed BLASTp (v2.12.0+) searches for all 71,528 sequences against the NCBI non-redundant (nr) protein database, with the Tardigrada taxonomy ID excluded to prevent self-hits. The search was focused on identifying homologs in closely related phyla such as Arthropoda and Nematoda. Sequences showing significant similarity to any non-tardigrade eukaryotic protein (E-value < 1e-3) were filtered out.

The remaining sequences were then subjected to a second filtering step to mitigate potential contamination from symbiotic or environmental microorganisms. This involved searches against a custom database of selected microbial proteomes extracted from UniProt. Specifically, based on prior studies of tardigrade microbiomes^46,47^, we screened against proteomes from *Cyanobacteria*, *Rickettsiales*, and *Polynucleobacter*. Sequences showing a moderate degree of similarity to microbial proteins (E-value < 1e-3) were considered potential contaminants and were excluded from the dataset. This stringent, multi-step curation process yielded a final dataset of 26,242 proteins, hereafter referred to as the tardigrade-specific proteome, which formed the basis for all subsequent discovery and analysis pipelines.

### Protein Clustering and Family Identification

To organize the curated proteome into putative protein families and to enable the discovery of novel ones, the 26,242 tardigrade-specific proteins were clustered based on sequence similarity using the CD-HIT algorithm^29^. Clustering was performed at a sequence identity threshold of 50% with a word size of three. This relatively permissive identity threshold was chosen to group potentially divergent homologs into broader, superfamily-level clusters, an approach well-suited for identifying novel families among rapidly evolving proteins such as IDPs, as recommended by the official user’s guide. The resulting clusters were analyzed to identify large, unannotated protein families. To evaluate the biological relevance of our approach, we then conducted keyword searches for ‘CAHS,’ ‘SAHS,’ ‘MAHS,’ and ‘Dsup’ within the annotations of all clustered proteins. This validation confirmed that our method successfully grouped members of known tardigrade-specific protein families into several discrete clusters, conferring high confidence on the newly identified, unannotated clusters as bona fide novel protein families.

### Acquisition of Sequences of Intrinsically Disordered Proteins and Structured Proteins for IDP Prediction Model

Sequences of intrinsically disordered proteins were obtained from the MobiDB database^48^ to construct the positive dataset for training the IDP prediction model. Specifically, we obtained two sequence datasets, one containing only sequences of proteins which have been experimentally validated to be intrinsically disordered, and another dataset containing sequences of proteins homologous to the experimentally validated proteins. The experimentally validated protein sequence dataset consists of 3,512 proteins and the dataset of proteins homologous to the experimentally validated proteins contains 374,919 proteins. To obtain sequences of structured proteins for the negative training set, the UniProt Knowledgebase (UniProtKB)^49^ was queried using a stringent set of criteria. Specifically, any proteins containing ‘disordered’ region annotations were excluded, and proteins which have a registered structure in the Protein Databank (PDB)^50^ or have an AlphaFold DB entry accompanied by domain information from the Pfam database^51^ was queried. To ensure high data quality, only manually reviewed proteins were included. This rigorous curation yielded a set of 422,137 structured proteins. These data were randomly divided into training and validation data at a 4:1 ratio and were used to train the IDP classifier.

### Machine Learning Model for Stress-Resistance Prediction

To prioritize candidate proteins for functional analysis, we developed AEGIS, a supervised machine learning classifier designed to predict a ‘stress-resistance potential’ score (AEGIS score). Recognizing that many known stress-resistance proteins are intrinsically disordered proteins (IDPs), we trained an IDP classifier for this purpose. Our framework began with feature engineering, converting each protein sequence into a high-dimensional embedding vector. We utilized ESM-C 300m to obtain embeddings from amino acid sequences^34^. To specialize these general-purpose protein embeddings for IDP features, Low-Rank Adaptation (LoRA)^35^ was employed. LoRA, with a rank of 4, an alpha of 16, and a dropout rate of 0.05, was applied to the ‘query,’ ‘key,’ and ‘value’ transformation layers within the attention structure of ESM-C 300m. The final feature representation for IDP prediction was a concatenation of the 960-dimensional CLS token embedding and the average of all amino acid residue embeddings. These combined features were then fed into a linear layer to predict IDP likelihood. During training, the model was optimized using the AdamW optimizer^52^. A linear warmup and linear decay schedule was employed with the initial learning rate of 5e-5. Batch size was set to 16, and the Binary Cross-Entropy was utilized for loss function. The model was trained for 7 epochs. The normalized logits from the trained model were then used as the AEGIS score. The Hugging-Face Transformers and PEFT (Parameter-Efficient Fine-Tuning) library (version 4.48.1, 0.15.2, respectively) were used to implement LoRA.

### IDR Prediction Model Architecture

To enable the mapping of IDRs of candidate proteins at residue-level resolution, a deep learning model which leverages residue-level protein embeddings to predict whether each residue of a protein belongs to an IDR was developed. Residue-level protein embeddings of dimension 960, are initially projected to a 1920-dimensional space via a linear layer. The projected embeddings are subsequently processed by multiple bi-directional LSTM layers, which process the input sequence in both the forward and reverse directions to effectively capture the intricate sequence dependencies among residues belonging to IDRs. Residual connections, layer normalizations and dropout are incorporated between the LSTM layers to ensure stable training and to prevent overfitting. The LSTM output is then fed into an enhanced positional encoding layer, which combines a fixed sinusoidal positional encoding (PE) with a learnable positional embedding (LPE), weighted by a trainable parameter α to incorporate positional information:

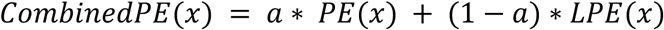

The encoded features are further fed into multiple Transformer encoder layers. Each Transformer layer comprises: a multi-head self-attention sub-layer, and a two-layer feed-forward network. Each of these sub-layers are accompanied by a residual connection step and a layer normalization step. Specifically, the multi-head self-attention sub-layer enables the model to attend to different parts of the sequence simultaneously. The attention scores are computed as:

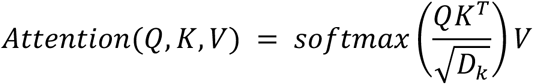

where *Q,, K*, and *V* are query, key and value matrices, and *D_k_* is the dimension of the key vector. A two-layer feed-forward network sub-layer with GELU activation and dropout expands dimensionality by a factor of 2 before projecting back. Layer normalization and residual connection is applied after each sub-layer. For a sub-layer function *Sublayer* the output is:

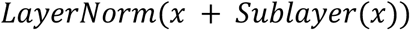

Learnable scaling factors (*λ_attn_* and *λ_FFN_*, initialized to 1 ∗ 10^-4^ ) are applied to the outputs of both the self-attention and feed-forward network sub-layers for training stability. The output following self-attention and residual connection becomes:

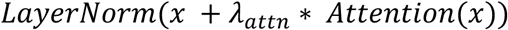

The output of the Transformer layers undergoes a final layer normalization and dropout. The feature is then fed into a classifier consisting of two linear layers with GELU activation and dropout, outputting probabilities for each residue being either disordered or structured.

### Data Preparation, Training and Optimization of IDR Prediction Model

Sequences of IDPs sourced from MobiDB^48^ and CAID^44^ were used for training and testing the IDR prediction model, respectively. Any protein sequences belonging to the CAID dataset were removed from the training dataset. Protein sequences exceeding a length of 1,000 amino acids were removed from the training dataset to reduce computation cost. This filtering process left 2,438 sequences, of which 80% was used as the training set, and 20% as the validation set. The protein sequences were embedded at the residue-level using the pre-trained ESM-C 300m model. For each training batch, consisting of multiple residue-level sequence embeddings and their labels, a custom collate function was implemented for batching, sorting sequences by length, and padding them to the maximum sequence length within each batch to enable efficient processing. AEGIS was trained using focal loss^53^ to address class imbalance, emphasizing hard-to-classify examples. Focal loss is defined as:

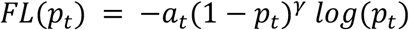

where *P_t_* is the predicted probability for the true class, *a_t_* is a weighting factor, and M is the focusing parameter. The AdamW^52^ optimizer was used for optimization. AEGIS was defined and trained using the PyTorch library (v2.5.1).

Hyperparameter optimization was performed using Optuna^54^ over 100 trials, with the objective set to maximize the F1-score on the validation set. Hyperparameters tuned included learning rate, hidden dimension, number of attention heads, number of LSTM and Transformer layers, dropout rate, forward expansion factor, scheduler type, weight decay and scheduler-specific parameters. Pruning was implemented using the median pruner to stop unpromising trials at an early stage. The exact search range and the optimal value of each hyperparameter is listed in Supplementary Table S3, S4, respectively. Each trial ran for 25 epochs. The best hyperparameters found with Optuna were then used for the final model evaluation. Model performance was assessed on the test set consisting solely of CAID protein sequences using area under the receiver operating characteristic curve (AU-ROC), Matthews Correlation Coefficient (MCC), and Fmax score and balanced accuracy (BACC).

### Integrative *In Silico* Characterization of Candidate Effectors

All tardigrade-specific proteins were scored using AEGIS. Subsequently, these proteins were embedded using the LoRA fine-tuned ESM-C 300m model. The high-dimensional embeddings were then reduced to lower dimensions using UMAP^55^, and clustered using the Leiden algorithm^56^ (with n_neighbors set to 500).

### Subcellular Localization Prediction

Subcellular localization was predicted using DeepLoc2.0^45^, with ESM1b specified as the backbone model for high-throughput analysis. This model calculates scores for ten subcellular compartments—nucleus, cytoplasm, extracellular, mitochondrion, cell membrane, endoplasmic reticulum, chloroplast, Golgi apparatus, lysosome/vacuole, and peroxisome—and outputs the localization with the highest score as the prediction. First, we validated the model’s prediction accuracy on known tardigrade disordered proteins (TDPs). Amino acid sequences of proteins registered as CAHS, MAHS, SAHS, and Dsup were retrieved from the UniProt database^49^. To avoid overlap with the DeepLoc2.0 training data, proteins with Gene Ontology (GO) annotations related to subcellular localization were excluded from our validation dataset. After this filtering, the validation set comprised 35 CAHS and 7 SAHS proteins (no proteins for MAHS or Dsup met the criteria). Based on previous research^9,10,16^, the primary localizations of CAHS, SAHS, MAHS, and Dsup were assumed to be the cytoplasm, extracellular space, mitochondrion, and nucleus, respectively. We then evaluated the concordance rate between the model’s predictions and these assumed localizations. Furthermore, the same localization analysis was applied to a set of TDP-like proteins identified from the *Ramazzottius varieornatus* proteome using PSI-BLAST (v2.12.0+)^57^, and to a subset of those proteins exhibiting particularly high AEGIS scores (Fig. 4c).

### Protein Structure Prediction

The tertiary structures of tardigrade-derived proteins were generated using the AlphaFold 3 (v3.0.1)^58^ prediction pipeline. The amino acid sequence of each target protein served as the initial input. A data preprocessing pipeline was then executed to construct a multiple sequence alignment (MSA) and search for structural templates. This process referenced the May 2025 release of the ‘alphafold3_database’. Subsequently, the structure prediction was calculated based on the sequence and structural information obtained from preprocessing, applying the official model parameters provided by Google DeepMind. To ensure reproducibility, the model’s random seed was fixed to 1, generating a single structural model for each target. All computations were performed in a GPU environment equipped with CUDA v12.8.0.

## ACKNOWLEDGEMENTS

This work was supported by KAKENHI grants from the Japan Society for the Promotion of Science (JSPS) to H.S. (24H01755 and 25H01571), as well as Takeda Science Foundation. We thank all the laboratory members for discussion and K. Tanaka for help with preparation of the manuscript.

## AUTHOR CONTRIBUTIONS

H.S. conceived of, designed, and supervised the study. T.I., K.O., and H.H. performed all formal analyses. T.I., K.O., H.H. and H.S. jointly wrote the manuscript. All authors have read and approved the final manuscript.

