## Supplementary Tables S1-S4, Supplementary Figure S1 for "A Functional Atlas of the Tardigrade Resistome Reveals a Diverse Molecular Toolkit for Extremotolerance"

#### Table S1: Counts of Tardigrade Protein Database Entries by Species

| # | Tardigrade species | Protein counts |
|---|---|---|
| 1 | Paramacrobiotus_metropolitanus | 25,574 |
| 2 | Ramazzottius_varieornatus | 22,844 |
| 3 | Hypsibius_exemplaris | 20,820 |
| 4 | Echiniscoides_sp. | 177 |
| 5 | Macrobiotus_sp. | 141 |
| 6 | Milnesium_sp. | 130 |
| 7 | Milnesium_tardigradum | 76 |
| 8 | Echiniscus_blumi | 74 |
| 9 | Ramazzottius_sp. | 66 |
| 10 | Acutuncus_antarcticus | 58 |
| 11 | Thulinius_stephaniae | 49 |
| 12 | Echiniscus_testudo | 48 |
| 13 | Ramazzottius_claudii | 48 |
| 14 | Paramacrobiotus_richtersi | 43 |
| 15 | Pseudechiniscus_sp. | 43 |
| 16 | Echiniscus_sp. | 32 |
| 17 | Mesobiotus_sp. | 32 |
| 18 | Rotaria_tardigrada | 27 |
| 19 | Viridiscus_sp. | 24 |
| 20 | Paramacrobiotus_sp. | 24 |
| 21 | Claxtonia_wendti | 23 |
| 22 | Macrobiotus_hufelandi | 22 |
| 23 | Xerobiotus_sp. | 22 |
| 24 | Paramacrobiotus_areolatus | 21 |
| 25 | Paramacrobiotus_fairbanksi | 19 |
| 26 | Parachela_sp. | 19 |
| 27 | Diphascon_sp. | 18 |
| 28 | Richtersius_sp. | 18 |
| 29 | Ramazzottius_oberhaeuseri | 15 |
| 30 | Cornechiniscus_lobatus | 15 |
| 31 | Paramacrobiotus_spOX=2719578 | 14 |
| 32 | Thulinius_sp. | 14 |
| 33 | Hypsibiidae_sp. | 14 |
| 34 | Batillipes_longispinosus | 13 |
| 35 | Hypsibius_sp. | 13 |
| 36 | Minibiotus_sp. | 12 |
| 37 | Macrobiotus_macrocalix | 12 |
| 38 | Macrobiotus_euxinus | 12 |
| 39 | Pseudobiotus_spinifer | 12 |
| 40 | Richtersius_coronifer | 11 |
| 41 | Claxtonia_sp. | 10 |
| 42 | Echiniscus_lineatus | 10 |
| 43 | Macrobiotus_polonicus | 10 |
| 44 | Hypsibius_dujardini | 10 |
| 45 | Macrobiotus_pallarii | 10 |
| 46 | Cryobiotus_klebelsbergi | 10 |
| 47 | Diaforobiotus_sp. | 9 |
| 48 | Acutuncus_sp. | 9 |
| 49 | Mesobiotus_furciger | 9 |
| 50 | Viridiscus_perviridis | 9 |
| 51 | Echiniscus_viridissimus | 8 |
| 52 | Bryodelphax_sp. | 8 |
| 53 | Actinarctus_doryphorus | 8 |
| 54 | Macrobiotus_joannae | 8 |
| 55 | Milnesium_tardigradumtardigradum | 7 |
| 56 | Diploechiniscus_oihonnae | 7 |
| 57 | Echiniscus_quadrispinosus | 7 |
| 58 | Macrobiotus_dolosus | 7 |
| 59 | Batillipes_sp. | 7 |
| 60 | Pilatobius_sp. | 7 |

| # | Tardigrade species | Protein counts |
|---|---|---|
| 61 | Macrobiotus_vladimiri | 7 |
| 62 | Macrobiotus_ripperi | 6 |
| 63 | Hypsibius_pallidoides | 6 |
| 64 | Diploechiniscus_sp. | 6 |
| 65 | Xerobiotus_reductus | 6 |
| 66 | Acutuncus_giovanniniae | 6 |
| 67 | Adropion_sp. | 6 |
| 68 | Xerobiotus_arenosum | 6 |
| 69 | Macrobiotus_wandae | 6 |
| 70 | Milnesium_inceptum | 6 |
| 71 | Platicrista_sp. | 6 |
| 72 | Crenubiotus_sp. | 6 |
| 73 | Diaforobiotus_islandicus | 6 |
| 74 | Hypsibius_scabropygus | 6 |
| 75 | Dactylobiotus_sp. | 6 |
| 76 | Dactylobiotus_parthenogeneticus | 6 |
| 77 | Echiniscus_tristis | 5 |
| 78 | Macrobiotus_kyoukenus | 5 |
| 79 | Echiniscus_canadensis | 5 |
| 80 | Echiniscus_spinulosus | 5 |
| 81 | Echiniscus_masculinus | 5 |
| 82 | Milnesium_berladnicorum | 5 |
| 83 | Macrobiotus_shonaicus | 5 |
| 84 | Astatumen_sp. | 5 |
| 85 | Echiniscus_merokensis | 5 |
| 86 | Echiniscus_granulatus | 5 |
| 87 | Mesobiotus_philippinicus | 5 |
| 88 | Macrobiotus_terminalis | 5 |
| 89 | Echiniscoides_sigismundi | 5 |
| 90 | Echiniscus_perarmatus | 5 |

| # | Tardigrade species | Protein counts |
|---|---|---|
| 91 | Hebesuncus_ryani | 4 |
| 92 | Pseudechiniscus_angelusalas | 4 |
| 93 | Mesobiotus_radiatus | 4 |
| 94 | Paramacrobiotus_depressus | 4 |
| 95 | Kristenseniscus_sp. | 4 |
| 96 | Mesocrista_revelata | 4 |
| 97 | Macrobiotus_siderophilus | 4 |
| 98 | Macrobiotus_kristenseni | 4 |
| 99 | Tenuibiotus_sp. | 4 |
| 100 | Xerobiotus_gretae | 4 |
| 101 | Dactylobiotus_spOX=2864188 | 4 |
| 102 | Echiniscoides_hoepneri | 4 |
| 103 | Murrayon_sp. | 4 |
| 104 | Cryoconicus_antiarktos | 3 |
| 105 | Macrobiotus_sandrae | 3 |
| 106 | Diphascon_puniceum | 3 |
| 107 | Minibiotus_furcatus | 3 |
| 108 | Hypsibioidea_sp. | 3 |
| 109 | Cornechiniscus_cornutus | 3 |
| 110 | Ursulinius_elegans | 3 |
| 111 | Barbaria_bigranulata | 3 |
| 112 | Hypsibius_pallidus | 3 |
| 113 | Tenuibiotus_voronkovi | 3 |
| 114 | Grevenius_sp. | 3 |
| 115 | Pilatobius_oculatus | 3 |
| 116 | Paramacrobiotus_spatialis | 3 |
| 117 | Echiniscus_scabrospinosus | 3 |
| 118 | Cornechiniscus_madagascariensis | 3 |
| 119 | Milnesium_eurystomum | 3 |
| 120 | Sisubiotus_spectabilis | 3 |

| # | Tardigrade species | Protein counts |
|---|---|---|
| 121 | Cornechiniscus_imperfectus | 3 |
| 122 | Macrobiotus_scoticus | 3 |
| 123 | Pseudechiniscus_facettalis | 3 |
| 124 | Macrobiotus_pseudopallarii | 3 |
| 125 | Mesobiotus_harmsworthi | 3 |
| 126 | Crenubiotus_ruhesteini | 3 |
| 127 | Bryodelphax_parvulus | 3 |
| 128 | Richtersius_ziemowiti | 3 |
| 129 | Claxtonia_mauccii | 3 |
| 130 | Macrobiotus_hannae | 3 |
| 131 | Paramurrayon_meieri | 3 |
| 132 | Mesobiotus_aradasi | 3 |
| 133 | Adropion_scoticum | 3 |
| 134 | Milnesium_pacificum | 3 |
| 135 | Nebularmis_reticulatus | 2 |
| 136 | Platicrista_angustata | 2 |
| 137 | Grevenius_granulifer | 2 |
| 138 | Mesobiotus_peterseni | 2 |
| 139 | Xerobiotus_naginae | 2 |
| 140 | Dactylobiotus_grandipes | 2 |
| 141 | Xerobiotus_pseudohufelandi | 2 |
| 142 | Barbaria_sp. | 2 |
| 143 | Macrobiotus_spOX=42245 | 2 |
| 144 | Echiniscus_spiniger | 2 |
| 145 | Adorybiotus_sp. | 2 |
| 146 | Echiniscus_trisetosus | 2 |
| 147 | Paramacrobiotus_arduus | 2 |
| 148 | Paramacrobiotus_celsus | 2 |
| 149 | Diphascon_pingue | 2 |
| 150 | Bertolanius_volubilis | 2 |

| # | Tardigrade species | Protein counts |
|---|---|---|
| 151 | Hypechiniscus_exarmatus | 2 |
| 152 | Paramacrobiotus_filipi | 2 |
| 153 | Claxtonia_molluscorum | 2 |
| 154 | Echiniscus_draconis | 2 |
| 155 | Kristenseniscus_limai | 2 |
| 156 | Ramazzottius_subanomalus | 2 |
| 157 | Crenubiotus_salishani | 2 |
| 158 | Minibiotus_sidereus | 2 |
| 159 | Isoechiniscoides_sifae | 2 |
| 160 | Macrobiotus_engbergi | 2 |
| 161 | Paramacrobiotus_experimentalis | 2 |
| 162 | Pilatobius_recamieri | 2 |
| 163 | Mesobiotus_huecoensis | 2 |
| 164 | Macrobiotus_annewintersae | 2 |
| 165 | Macrobiotus_margoae | 2 |
| 166 | Macrobiotus_mileri | 2 |
| 167 | Echiniscus_pellucidus | 2 |
| 168 | Milnesium_reductum | 2 |
| 169 | Macrobiotus_polypiformis | 2 |
| 170 | Milnesium_lagniappe | 2 |
| 171 | Sisubiotus_hakaiensis | 2 |
| 172 | Ramazzottius_szeptycki | 2 |
| 173 | Paramacrobiotus_gadabouti | 2 |
| 174 | Halobiotus_crispae | 2 |
| 175 | Macrobiotus_fontourai | 2 |
| 176 | Milnesium_dornensis | 2 |
| 177 | Minibiotus_citlalium | 2 |
| 178 | Pseudobiotus_sp. | 2 |
| 179 | Batillipes_pennaki | 2 |
| 180 | Milnesium_pentapapillatum | 2 |

| # | Tardigrade species | Protein counts |
|---|---|---|
| 181 | Viridiscus_viridissimus | 2 |
| 182 | Diaforobiotus_hyperonyx | 2 |
| 183 | Echiniscus_insularis | 2 |
| 184 | Nebularmis_japonicus | 2 |
| 185 | Tardigrada_sp. | 2 |
| 186 | Echiniscus_latruncularis | 2 |
| 187 | Testechiniscus_spitsbergensis | 2 |
| 188 | Mesobiotus_occultatus | 2 |
| 189 | Cornechiniscus_subcornutus | 2 |
| 190 | Echiniscus_manuelae | 2 |
| 191 | Hebesuncus_conjugens | 2 |
| 192 | Echiniscus_aonikenk | 1 |
| 193 | Tenuibiotus_danilovi | 1 |
| 194 | Echiniscus_tantulus | 1 |
| 195 | Echiniscus_belloporus | 1 |
| 196 | Echiniscus_siticulosus | 1 |
| 197 | Mesocrista_sp. | 1 |
| 198 | Barbaria_ganczareki | 1 |
| 199 | Adorybiotus_granulatus | 1 |
| 200 | Testechiniscus_laterculus | 1 |
| 201 | Echiniscus_robertsi | 1 |
| 202 | Ramazzottius_groenlandensis | 1 |
| 203 | Halechiniscus_sp. | 1 |
| 204 | Tenuibiotus_tenuiformis | 1 |
| 205 | Macrobiotus_glebkai | 1 |
| 206 | Macrobiotus_kamilae | 1 |
| 207 | Dactylobiotus_selenicus | 1 |
| 208 | Macrobiotus_sottilei | 1 |
| 209 | Milnesioides_sp. | 1 |
| 210 | Milnesium_rastrum | 1 |

| # | Tardigrade species | Protein counts |
|---|---|---|
| 211 | Echiniscus_meridionalis | 1 |
| 212 | Macrobiotus_kosmali | 1 |
| 213 | Acanthechiniscus_goedeni | 1 |
| 214 | Arctodiphascon_tenue | 1 |
| 215 | Macrobiotus_rybaki | 1 |
| 216 | Echiniscus_dentatus | 1 |
| 217 | Echiniscus_capensis | 1 |
| 218 | Echiniscus_attenboroughi | 1 |
| 219 | Ramazzottius_kretschmanni | 1 |
| 220 | Macrobiotus_crustulus | 1 |
| 221 | Mesobiotus_fiedleri | 1 |
| 222 | Minibiotus_pentannulatus | 1 |
| 223 | Milnesium_wrightae | 1 |
| 224 | Minibiotus_ioculator | 1 |
| 225 | Grevenius_asper | 1 |
| 226 | Mesocrista_spitzbergensis | 1 |
| 227 | Paramacrobiotus_lachowskae | 1 |
| 228 | Milnesium_matheusi | 1 |
| 229 | Echiniscus_similaris | 1 |
| 230 | Echiniscus_scabrocirrosus | 1 |
| 231 | Echiniscus_oreas | 1 |
| 232 | Echiniscus_lichenorum | 1 |
| 233 | Echiniscus_lapponicus | 1 |
| 234 | Echiniscus_intricatus | 1 |
| 235 | Echiniscus_imitans | 1 |
| 236 | Echiniscus_gracilis | 1 |
| 237 | Minibiotus_intermedius | 1 |
| 238 | Macrobiotus_ariekammensisgroenlandicus | 1 |
| 239 | Macrobiotus_kirghizicus | 1 |
| 240 | Macrobiotus_ariekammensisariekammensis | 1 |

| # | Tardigrade species | Protein counts |
| --- | --- | --- |
| 241 | Nebularmis_crebraclava | 1 |
| 242 | Echiniscus_hoonsooi | 1 |
| 243 | Echiniscus_clevelandi | 1 |
| 244 | Styraconyx_takeshii | 1 |
| 245 | Mesobiotus_marmoreus | 1 |
| 246 | Mesobiotus_imperialis | 1 |
| 247 | Mesobiotus_skorackii | 1 |
| 248 | Echiniscus_peruvianus | 1 |
| 249 | Echiniscus_evelinae | 1 |
| 250 | Milnesium_pseudotardigradum | 1 |
| 251 | Macrobiotus_almadai | 1 |
| 252 | Milnesium_almatyense | 1 |
| 253 | Nebularmis_indicus | 1 |
| 254 | Nebularmis_burmensis | 1 |
| 255 | Nebularmis_auratus | 1 |
| 256 | Echiniscus_charrua | 1 |
| 257 | Echiniscus_cavagnaroi | 1 |
| 258 | Barbaria_danieli | 1 |
| 259 | Barbaria_madonnae | 1 |
| 260 | Echiniscoides_pollocki | 1 |
| 261 | Pseudechiniscus_suillus | 1 |
| 262 | Pseudechiniscus_indistinctus | 1 |
| 263 | Pseudechiniscus_ehrenbergi | 1 |
| 264 | Pseudechiniscus_lacyformis | 1 |
| 265 | Pseudechiniscus_juanitae | 1 |
| 266 | Barbaria_ollantaytamboensis | 1 |
| 267 | Cornechiniscus_holmeni | 1 |
| 268 | Viridiscus_viridianus | 1 |
| 269 | Bertolanius_weglarskae | 1 |
| 270 | Isohypsibius_prosostomus | 1 |

| # | Tardigrade species | Protein counts |
| --- | --- | --- |
| 271 | Dastychius_improvisus | 1 |
| 272 | Pilatobius_islandicus | 1 |
| 273 | Paramacrobiotus_tonollii | 1 |
| 274 | Acutuncus_mecnuffi | 1 |
| 275 | Macrobiotus_occidentalis | 1 |
| 276 | Borealibius_zetlandicus | 1 |
| 277 | Mesobiotus_diegoi | 1 |
| 278 | Mesobiotus_maklowiczi | 1 |
| 279 | Macrobiotus_paulinae | 1 |
| 280 | Florarctus_sp. | 1 |
| 281 | Milnesiidae_sp. | 1 |
| 282 | Macrobiotidae_sp. | 1 |
| 283 | Isohypsibius_sp. | 1 |
| 284 | Parechiniscus_chitonides | 1 |
| 285 | Proechiniscus_hanneae | 1 |
| 286 | Antechiniscus_lateromamillatus | 1 |
| 287 | Pseudechiniscus_islandicus | 1 |
| 288 | Pseudechiniscus_novaezelandiae | 1 |
| 289 | Doryphoribius_sp. | 1 |
| 290 | Hypechiniscus_gladiator | 1 |
| 291 | Mopsechiniscus_granulosus | 1 |
| 292 | Ramajendas_frigidus | 1 |
| 293 | Cryoconicus_kaczmareki | 1 |
| 294 | Ramazzottius_rupeus | 1 |
| 295 | Thulinius_augusti | 1 |
| 296 | Mesobiotus_romani | 1 |
| 297 | Astatumen_trinacriae | 1 |
| 298 | Mesobiotus_datanlanicus | 1 |
| 299 | Hypsibius_convergens | 1 |
| 300 | Mesobiotus_hilariae | 1 |

| # | Tardigrade species | Protein counts |
|---|---|---|
| 301 | Mesobiotus_dilimanensis | 1 |
| 302 | Dactylobiotus_ovimutans | 1 |
| 303 | Grevenius_pushkini | 1 |
| 304 | Diphascon_higginsi | 1 |
| 305 | Thulinius_ruffoi | 1 |
| 306 | Macrobiotus_canaricus | 1 |
| 307 | Macrobiotus_papei | 1 |
| 308 | Richtersius_tertius | 1 |
| 309 | Minibiotus_gumersindoi | 1 |
| 310 | Cornechiniscus_sp. | 1 |
| 311 | Pseudechiniscus_novaezeelandiae | 1 |
| 312 | Phallocephale_tallagandensis | 1 |
| 313 | Mesobiotus_silesiacus | 1 |
| 314 | Novechiniscus_armadilloides | 1 |
| 315 | Cucumibius_annulatus | 1 |
| 316 | Pilatobius_bullatus | 1 |
| 317 | Adropion_fagineum | 1 |
| 318 | Minibiotus_dispositus | 1 |
| 319 | Pseudechiniscus_shintai | 1 |
| 320 | Echiniscus_ornamentatus | 1 |
| 321 | Echiniscus_succineus | 1 |
| 322 | Milnesium_alpigenum | 1 |
| 323 | Barbaria_jenningsi | 1 |
| 324 | Echiniscus_mediantus | 1 |
| 325 | Mesobiotus_insanis | 1 |
| 326 | Macrobiotus_caelestis | 1 |
| 327 | Tenuibiotus_zandrae | 1 |
| 328 | Eremobiotus_alicatai | 1 |
| 329 | Murrayon_dianeae | 1 |
| 330 | Murrayon_pullari | 1 |
| 331 | Mesobiotus_emiliae | 1 |
| 332 | Xerobiotus_inermis | 1 |
| 333 | Pseudohexapodibius_degenerans | 1 |
| 334 | Xerobiotus_litus | 1 |
| 335 | Echiniscus_pusae | 1 |
| 336 | Echiniscus_minutus | 1 |
| 337 | Echiniscus_africanus | 1 |
| 338 | Macrobiotus_ovovittatus | 1 |
| 339 | Calohypsibius_ornatus | 1 |
| 340 | Raribius_sp. | 1 |
| 341 | Platicrista_horribilis | 1 |
| 342 | Guidettion_prorsirostre | 1 |
| 343 | Itaquascon_sp. | 1 |
| 344 | Astatumen_bartosi | 1 |
| 345 | Macrobiotus_rebecchii | 1 |
| 346 | Macrobiotus_birendrai | 1 |
| 347 | Echiniscoides_rugostellatus | 1 |
| 348 | Echiniscus_virginicus | 1 |
| 349 | Pseudechiniscus_dastychi | 1 |
| 350 | Pseudechiniscus_quadrilobatus | 1 |
| 351 | Mesobiotus_ethiopicus | 1 |
| 352 | Milnesium_variefidum | 1 |
| 353 | Macrobiotus_noongaris | 1 |
| 354 | Diaforobiotus_svalbardicus | 1 |
| 355 | Crenubiotus_crenulatus | 1 |
| 356 | Unclassified | 108 |
|  | Total | 71,528 |

#### Table S2: Summary of Tardigrade-Specific Protein Clusters (Size ≥15)

| Cluster no. | Representative Protein ID | Representative Protein Name | Cluster Size | Predicted Localization | Mean AEGIS score |
|---|---|---|---|---|---|
| 1 | A0A1D1UX84_RAMVA | AMOP domain-containing protein | 67 | Extracellular | 0.475 |
| 2 | XP_055357560 | rootletin-like | 64 | Cytoplasm | 0.737 |
| 3 | A0A1D1V3I0_RAMVA | Spore coat protein | 58 | Extracellular | 0.548 |
| 4 | XP_055357440 | uncharacterized protein LOC129602448 isoform X2 | 57 | Cytoplasm | 0.431 |
| 5 | XP_055356973 | RNA-binding protein 25-like | 53 | Nucleus | 0.811 |
| 6 | XP_055357138 | uncharacterized protein LOC129602190 | 48 | Extracellular | 0.534 |
| 7 | CAHS2_HYPEX | Cytosolic-abundant heat soluble protein 86272 | 44 | Cytoplasm | 0.676 |
| 8 | XP_055357422 | uncharacterized protein LOC129602431 | 37 | Cytoplasm\|Nucleus | 0.496 |
| 9 | A0A1D1UMT1_RAMVA | Major facilitator superfamily (MFS) profile domain-containing protein | 33 | Cytoplasm | 0.677 |
| 10 | XP_055357072 | uncharacterized protein LOC129602125 | 32 | Extracellular | 0.447 |
| 11 | CAHS8_HYPEX | Cytosolic-abundant heat soluble protein 94063 | 28 | Cytoplasm | 0.67 |
| 12 | XP_055354689 | uncharacterized protein LOC129600250 | 27 | Cytoplasm | 0.43 |
| 13 | A0A1D1W8E7_RAMVA | Reverse transcriptase domain-containing protein | 25 | Cytoplasm\|Nucleus | 0.608 |
| 14 | A0A1W0XBY0_HYPEX | Tissue-type plasminogen activator | 24 | Extracellular | 0.536 |
| 15 | XP_055355134 | uncharacterized protein LOC129600621 | 23 | Nucleus | 0.541 |
| 16 | A0A9X6NM28_HYPEX | CCHC-type domain-containing protein (Fragment) | 23 | Cytoplasm\|Nucleus | 0.704 |
| 17 | XP_055356946 | uncharacterized protein LOC129602030 isoform X2 | 23 | Nucleus | 0.581 |
| 18 | XP_055354526 | uncharacterized protein LOC129600139 | 22 | Cytoplasm | 0.757 |
| 19 | XP_055356238 | uncharacterized protein | 21 | Cytoplasm | 0.492 |

| | | | | | |
|---|---|---|---|---|---|
| | | LOC129601452 | | | |
| 20 | XP_055356306 | uncharacterized protein LOC129601495 | 21 | Extracellular | 0.383 |
| 21 | XP_055357444 | uncharacterized protein LOC129602451 | 21 | Cytoplasm\|Nucleus | 0.534 |
| 22 | SAHS2_HYPEX | Secretory-abundant heat soluble protein 53582 | 21 | Extracellular | 0.497 |
| 23 | XP_055345589 | loricrin-like | 21 | Extracellular | 0.53 |
| 24 | XP_055355952 | uncharacterized protein LOC129601222 | 20 | Cytoplasm | 0.51 |
| 25 | CAHS5_HYPEX | Cytosolic-abundant heat soluble protein 89226 | 20 | Cytoplasm | 0.65 |
| 26 | A0A1D1UNH4_RAMVA | RRM domain-containing protein | 20 | Cytoplasm\|Nucleus | 0.617 |
| 27 | A0A1W0X3P1_HYPEX | ABC transporter domain-containing protein | 19 | Cytoplasm | 0.656 |
| 28 | XP_055349496 | uncharacterized protein LOC129596284 | 19 | Endoplasmic reticulum / Lysosome / Vacuole | 0.568 |
| 29 | A0A1D1V5U5_RAMVA | Uncharacterized protein | 19 | Extracellular | 0.47 |
| 30 | WPK49534 | AHS alpha | 19 | Extracellular | 0.845 |
| 31 | XP_055357329 | putative leucine-rich repeat-containing protein DDB_G0290503 | 18 | Cytoplasm | 0.757 |
| 32 | XP_055357038 | uncharacterized protein LOC129602096 | 18 | Cytoplasm | 0.404 |
| 33 | XP_055357233 | uncharacterized protein LOC129602261 isoform X1 | 18 | Cytoplasm\|Nucleus | 0.543 |
| 34 | XP_055351037 | uncharacterized protein LOC129597490 isoform X2 | 18 | Extracellular | 0.475 |
| 35 | A0A1D1WBX2_RAMVA | Homeobox domain-containing protein | 17 | Nucleus | 0.544 |
| 36 | A0A1D1W6W4_RAMVA | Tudor-knot domain-containing protein | 17 | Nucleus | 0.693 |
| 37 | XP_055351512 | cadherin-related family member 5-like isoform X1 | 16 | Nucleus | 0.782 |
| 38 | XP_055357374 | uncharacterized protein LOC129602394 | 16 | Cytoplasm | 0.443 |
| 39 | A0A1D1VFW1_RAMVA | cGMP-dependent protein kinase N-terminal coiled-coil domain-containing protein | 16 | Nucleus | 0.714 |
| 40 | A0A1W0W8Q0_HYPEX | Replicase polyprotein 1a | 16 | Cytoplasm | 0.85 |

| | | | | | |
|---|---|---|---|---|---|
| 41 | XP_055357490 | uncharacterized protein LOC129602490 | 16 | Extracellular | 0.474 |
| 42 | WPK49552 | AHS beta, partial | 16 | Extracellular | 0.82 |
| 43 | XP_055355962 | caldesmon-like | 15 | Nucleus | 0.776 |
| 44 | XP_055357174 | uncharacterized protein LOC129602209 isoform X5 | 15 | Cell membrane | 0.72 |
| 45 | XP_055352113 | uncharacterized protein LOC129598300 isoform X2 | 15 | Cytoplasm | 0.416 |
| 46 | XP_055355862 | uncharacterized protein LOC129601154 | 15 | Extracellular | 0.41 |
| 47 | A0A1D1VPD8_RAMVA | Nose resistant-to-fluoxetine protein N-terminal domain-containing protein | 15 | Extracellular | 0.552 |
| 48 | A0A1D1VVS9_RAMVA | Uncharacterized protein | 15 | Nucleus | 0.675 |
| 49 | A0A1D1UY52_RAMVA | SMP domain-containing protein | 15 | Nucleus | 0.75 |
| 50 | A0A1D1VIJ0_RAMVA | SAM domain-containing protein | 15 | Nucleus | 0.725 |

**Table S3: Search Space of Hyperparameters Tuned using Optuna**

| Hyperparameter | Search Space / Range |
|---|---|
| Learning rate | Uniform(1e-5, 1e-3) |
| LSTM hidden dimension | Categorical([32, 64, 128, 256, 512]) |
| Number of attention heads | Categorical([1, 2, 4, 8]) |
| Number of LSTM layers | IntUniform(1, 3) |
| Number of Transformer layers | IntUniform(1, 4) |
| Dropout rate | Uniform(0.1, 0.5) |
| Forward expansion factor | Categorical([2, 4]) |
| Learning rate scheduler | Categorical([Plateau, Cosine]) |
| Weight decay | Uniform(1e-5, 1e-2) |
| Scheduler patience | IntUniform(3, 10) |

**Table S4: Optimal Hyperparameters Discovered using Optuna**

| Hyperparameter | Optimal value |
|---|---|
| Learning rate | 3.4997e-05 |
| LSTM hidden dimension | 64 |
| Number of attention heads | 2 |
| Number of LSTM layers | 3 |
| Number of Transformer layers | 4 |
| Dropout rate | 0.4599 |
| Forward expansion factor | 2 |
| Learning rate scheduler | Plateau |
| Weight decay | 0.0073 |
| Scheduler patience | 10 |

### Supplementary Figure Legends

**Supplementary Figure 1 | Architecture of the AEGIS IDR Prediction Model**

The AEGIS IDR prediction model predicts disorder probabilities for each amino acid residue of an input protein sequence. An input protein sequence is first converted into residue-level embeddings using a pre-trained ESM-C 300m model. These residue-level embeddings are then processed by a stack of three layers of bi-directional Long Short-Term Memory (LSTM) layers to capture local sequence context. To incorporate positional information, a weighted positional embedding, calculated as a learnable combination of sinusoidal positional encoding and learned positional embedding is added to the output of the LSTM layers. The resulting vectors are then fed into four Transformer encoder layers, which model long-range dependencies. Each Transformer layer is composed of a multi-head self-attention and feed-forward network, with residual connections and layer normalization. A fully connected layer acts as the output head, predicting a disorder probability for each amino acid residue of the input sequence.

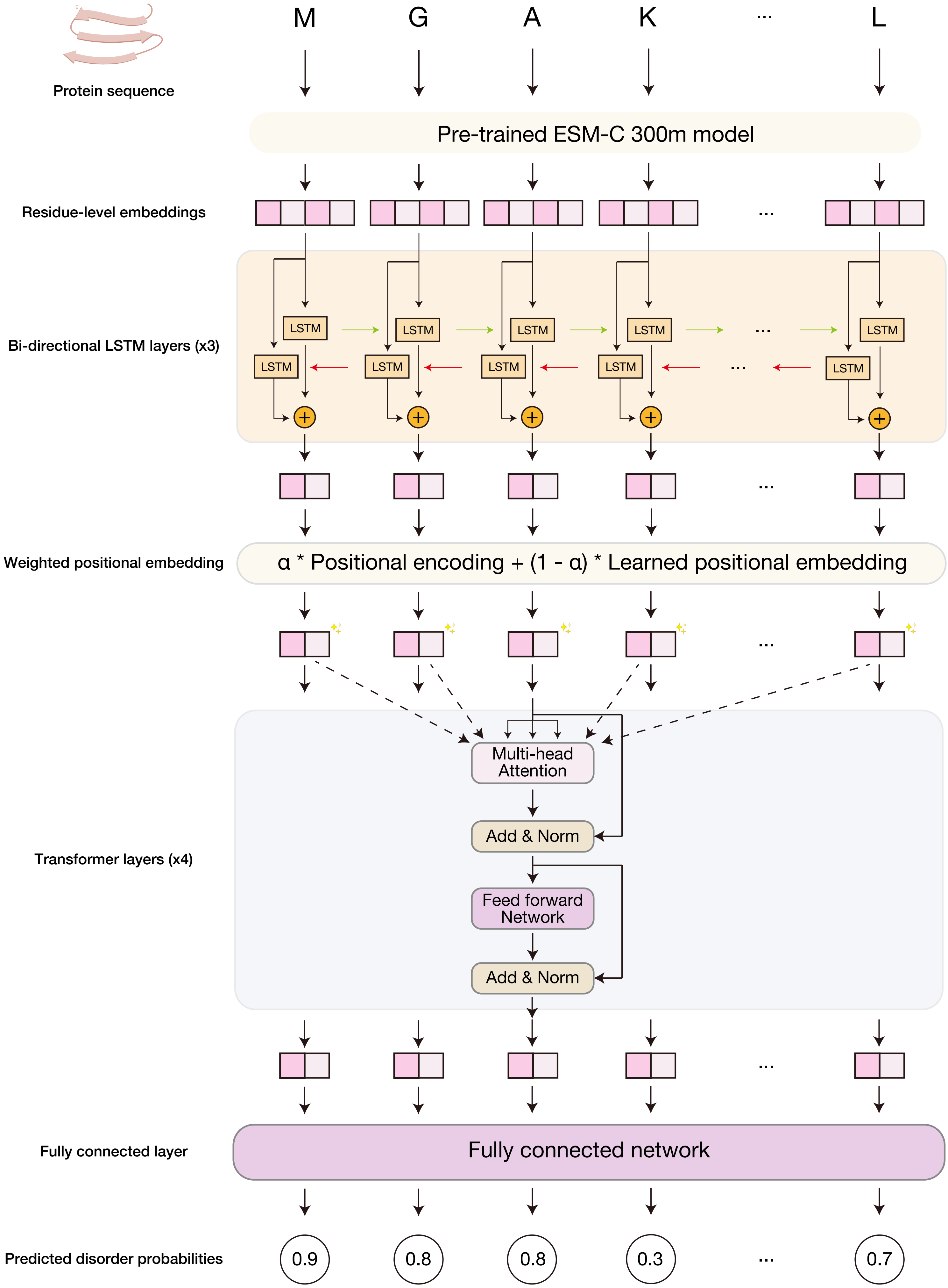


**Ito et al., Supplementary Figure 1**
